# Impaired T-cell and antibody immunity after COVID-19 infection in chronically immunosuppressed transplant recipients

**DOI:** 10.1101/2021.05.03.442371

**Authors:** Chethan Ashokkumar, Vinayak Rohan, Alexander H Kroemer, Sohail Rao, George Mazariegos, Brandon W Higgs, Satish Nadig, Jose Almeda, Harmeet Dhani, Khalid Khan, Nada Yazigi, Udeme Ekong, Stuart Kaufman, Monica M Betancourt-Garcia, Kavitha Mukund, Pradeep Sethi, Shikhar Mehrotra, Kyle Soltys, Manasi S Singh, Geoffrey Bond, Ajai Khanna, Mylarappa Ningappa, Brianna Spishock, Elizabeth Sindhi, Neha Atale, Maggie Saunders, Prabhakar Baliga, Thomas Fishbein, Shankar Subramaniam, Rakesh Sindhi

## Abstract

Assessment of T-cell immunity to the COVID-19 coronavirus requires reliable assays and is of great interest, given the uncertain longevity of the antibody response. Some recent reports have used immunodominant spike (S) antigenic peptides and anti-CD28 co-stimulation in varying combinations to assess T-cell immunity to SARS-CoV-2. These assays may cause T-cell hyperstimulation and could overestimate antiviral immunity in chronically immunosuppressed transplant recipients, who are predisposed to infections and vaccination failures. Here, we evaluate CD154-expressing T-cells induced by unselected S antigenic peptides in 204 subjects-103 COVID-19 patients and 101 healthy unexposed subjects. Subjects included 72 transplanted and 130 non-transplanted subjects. S-reactive CD154+T-cells co-express and can thus substitute for IFNγ (n=3). Assay reproducibility in a variety of conditions was acceptable with coefficient of variation of 2-10.6%. S-reactive CD154+T-cell frequencies were a) higher in 42 healthy unexposed transplant recipients who were sampled pre-pandemic, compared with 59 healthy non-transplanted subjects (p=0.02), b) lower in Tr COVID-19 patients compared with healthy transplant patients (p<0.0001), c) lower in Tr patients with severe COVID-19 (p<0.0001), or COVID-19 requiring hospitalization (p<0.05), compared with healthy Tr recipients. S-reactive T-cells were not significantly different between the various COVID-19 disease categories in NT recipients. Among transplant recipients with COVID-19, cytomegalovirus co-infection occurred in 34%; further, CMV-specific T-cells (p<0.001) and incidence of anti-receptor-binding-domain IgG (p=0.011) were lower compared with non-transplanted COVID-19 patients. Healthy unexposed transplant recipients exhibit pre-existing T-cell immunity to SARS-CoV-2. COVID-19 infection leads to impaired T-cell and antibody responses to SARS-CoV-2 and increased risk of CMV co-infection in transplant recipients.

## Introduction

In chronically immunosuppressed transplant recipients (Tr), the status of immunity to COVID-19 infection is of great interest. This population is prone to life-threatening consequences of viral infection and failure of vaccination during periods marked by use of high-dose immunosuppression^1^. Lifelong use of anti-rejection immunosuppressants contributes to this impairment and may also limit post-infectious and post-vaccination immunity to SARS-CoV-2 ^2,3^. Although antibodies can be demonstrated after natural COVID-19 infection and vaccination in the general population, this information is not available for Tr recipients^4-11^. Pre-existing T-cells that recognize SARS-CoV-2 are another component of immunity to this virus^12-15^. This type of immunity arises from prior exposure to human coronaviruses (hCoV), which account for 15% of seasonal flu and have structural similarities to SARS-CoV-2^16-17^. Pre-existing cellular immunity may also compensate for impaired antibody responses to COVID-19 infection and vaccination, and aid in combating variant strains that are starting to emerge. T-cell immunity may also reassure those individuals wishing to re-engage with the general public, but who are unable to tolerate vaccination or fail to achieve a durable antibody response. Pre-existing cellular immunity to SARS-CoV-2 has been demonstrated in non-transplanted subjects, but not in Tr recipients^12-15^.

Recently proposed assays which measure T-cell immunity to SARS-CoV-2 may need to be modified to characterize T-cell immunity in Tr recipients. Some assays stimulate T-cells with those peptides representing the spike protein S, which have high affinity to well represented HLA specificities in a given population^13,14^. Such peptide mixtures can potentially overstimulate T-cells from individuals with these HLA specificities, but not T-cells from underrepresented individuals. Other assays also use the co-stimulators, anti-CD28 alone, or with anti-CD49d ^12,14^. These adjunctive stimuli can also lead to an overestimate of T-cell immunity. Clinical decisions founded on such overestimates can be falsely reassuring in chronically immunosuppressed patients, and lead to errors in clinical judgement with adverse consequences. Some assays also use cytotoxic intracellular staining procedures, or only count those cells which co-express multiple markers as antigen-reactive. Such multi-marker assays require large numbers of cells from individuals with COVID-19 infection, who can be severely lymphopenic, and sophisticated laboratories. Another challenge is extrapolating findings from these studies, all of which have been performed in non-transplant (NT) subjects who were recently diagnosed or were convalescing, to Tr recipients. Higher frequencies of S-reactive T-cells were observed in convalescent and non-critically ill COVID-19 NT patients compared with healthy unexposed individuals. S-reactive frequencies were undetectable in critically ill COVID-19 patients^12-15^.

A clinically usable test design is exemplified by assays to measure T-cell response to cytomegalovirus (CMV). These assays use unselected peptide mixtures representing the entire antigenic sequence of interest, a single activation marker, and no costimulators^18-20^. Here, we describe a minimal marker assay to characterize S-reactive T- and B-cells in healthy unexposed subjects and COVID-19 patients most of whom were hospitalized, with an emphasis on chronically immunosuppressed solid organ transplant recipients. A sizeable cohort of NT subjects is also included to enable robust conclusions and comparisons.

## Results

Human Subjects: Of 204 total subjects, 101 were healthy subjects, H-Tr or H-NT, and 103 had been recently diagnosed with COVID-19, Tr or NT. The 204 subjects included 74 Tr recipients, of whom 42 were sampled pre-pandemic in 2019 or earlier, and 32 had COVID-19 infection. Of 130 NT subjects, 59 were H-NT subjects of whom twenty-five were sampled pre-pandemic and thirty-four were negative for COVID-19 by antibody testing. Seventy-one NT subjects had COVID-19 infection. Compared with healthy unexposed subjects, COVID-19 patients were predominantly non-Caucasian (38/101 vs 83/103 non-Caucasians, p<0.001) males (47/101 vs 60/103 males, p=NS) and were significantly older (41 vs 54 years, p=7.7E-06). General demographics for all 204 subjects are summarized in Table 1. Details including treatment and outcomes for COVID-19 patients are shown in Table S1.

**Table 1:**
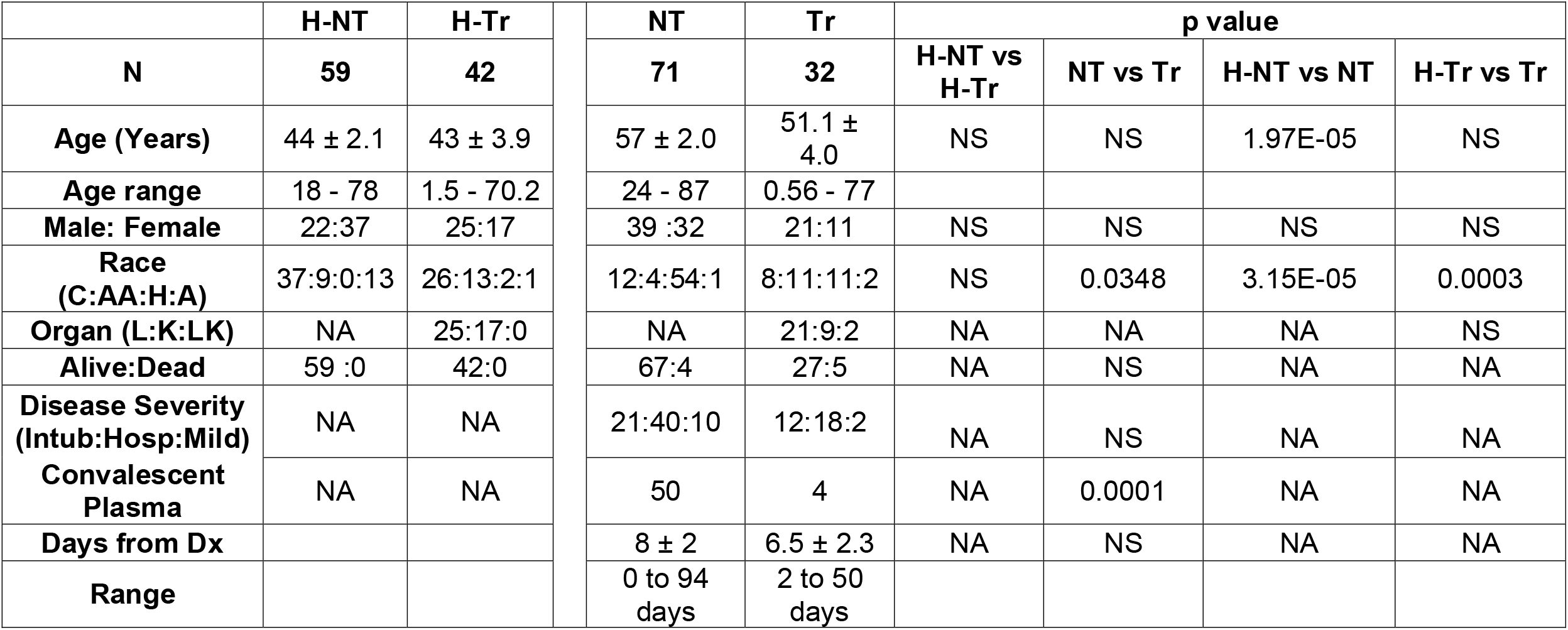
General demographics of the study population. Abbreviations: H-Tr: Healthy transplant, H-NT: healthy non-transplant, Tr-transplant recipients with COVID-19, NT-non-transplant patient with COVID-19, C: Caucasian, AA: African American, H: Hispanic, A: Asian, L: Liver transplant, K: Kidney transplant and LK: Liver-Kidney Transplant, Intub: Intubation, Hosp: Hospitalized, Mild: Mild.

|  | H-NT | H-Tr |  | NT | Tr | p value |  |  |  |
| --- | --- | --- | --- | --- | --- | --- | --- | --- | --- |
| N | 59 | 42 |  | 71 | 32 | H-NT vs H-Tr | NT vs Tr | H-NT vs NT | H-Tr vs Tr |
| Age (Years) | 44 ± 2.1 | 43 ± 3.9 |  | 57 ± 2.0 | 51.1 ± 4.0 | NS | NS | 1.97E-05 | NS |
| Age range | 18 - 78 | 1.5 - 70.2 |  | 24 - 87 | 0.56 - 77 |  |  |  |  |
| Male: Female | 22:37 | 25:17 |  | 39 :32 | 21:11 | NS | NS | NS | NS |
| Race (C:AA:H:A) | 37:9:0:13 | 26:13:2:1 |  | 12:4:54:1 | 8:11:11:2 | NS | 0.0348 | 3.15E-05 | 0.0003 |
| Organ (L:K:LK) | NA | 25:17:0 |  | NA | 21:9:2 | NA | NA | NA | NS |
| Alive:Dead | 59 :0 | 42:0 |  | 67:4 | 27:5 | NA | NS | NA | NA |
| Disease Severity (Intub:Hosp:Mild) | NA | NA |  | 21:40:10 | 12:18:2 | NA | NS | NA | NA |
| Convalescent Plasma | NA | NA |  | 50 | 4 | NA | 0.0001 | NA | NA |
| Days from Dx |  |  |  | 8 ± 2 | 6.5 ± 2.3 | NA | NS | NA | NA |
| Range |  |  |  | 0 to 94 days | 2 to 50 days |  |  |  |  |

### SARS2-specific CD154-expressing T- and B-cells co-express IFNγ and interleukin-6 (IL-6)

Stimulation of PBL from three healthy-NT with S peptides, cell permeabilization and ICS with specific fluorochrome-labeled detector antibodies showed that S-reactive T-cells which expressed CD154 also expressed IFNγ, a marker of cytotoxic T-cells (Figure 1a). Further, CD154 was also co-expressed with IL-6 in B-cells (Figure 1b). Thus, CD154 was used as a surrogate for S-reactive IFNγ+T-cells, and S-reactive IL-6+B-cells in all subsequent assays using nonpermeabilizing CD154 staining described previously^18,21^.

**Figure 1.**
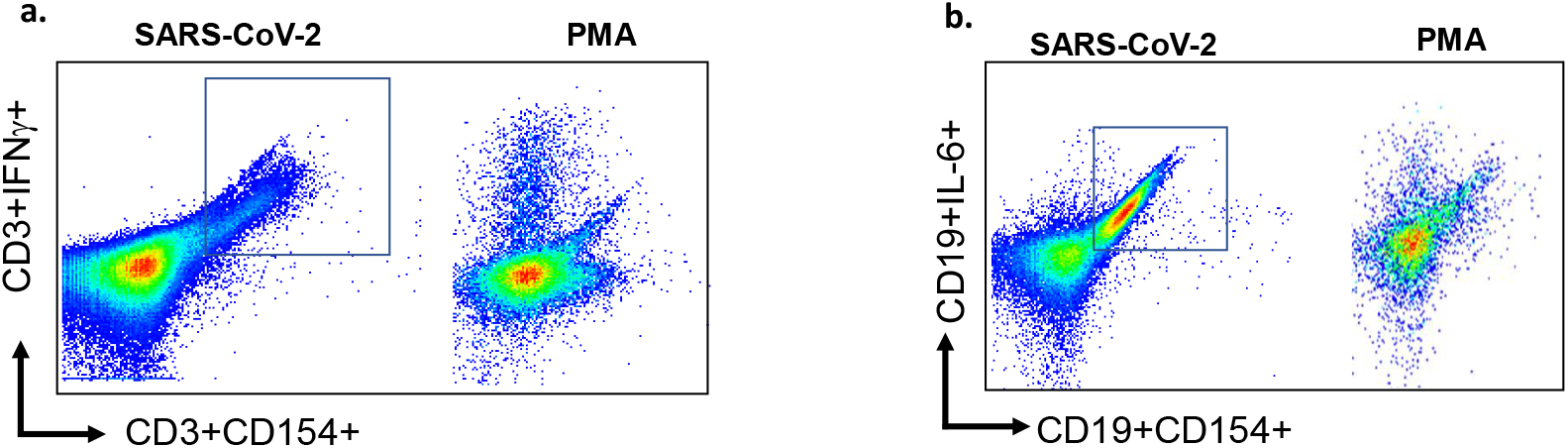
Flow cytometry scatterplots show a) expression of CD154 and IFNγ in S-reactive and PMA-reactive T-cells, and (b) expression of CD154 and IL-6 in S-reactive and PMA-reactive B-cells. PMA=phorbol-myristic acid, a mitogen.

### Reproducibility

S-reactive CD154-expressing CD3, CD4, CD8 and CD19 cells were measured in duplicate assays performed on the same day, before and after 7 days of cryopreservation in liquid nitrogen, and before and after overnight storage or overnight shipment at ambient temperature. Mean coefficient of variation between duplicate assays was 2-10.6% in these various conditions (Tables S2-S5).

### T- and B-cell responses to spike antigens are impaired with COVID-19 infection and increasing disease severity

Frequencies of S-reactive CD3, CD4, CD8 and CD19 B-cells were lower in 32 Tr recipients with COVID-19 compared with 42 H-Tr recipients (p<0.001) (Figure 2a-b, Table S6). S-reactive CD3 and S-reactive CD8 cell frequencies decreased progressively with increasing COVID-19 severity in Tr patients with COVID-19 infection. This decrease achieved significance for hospitalized recipients and those with severe COVID-19, compared with healthy-Tr subjects (Figure 2c-d). S-reactive T-cell frequencies in Tr patients with mild COVID-19 infection were similar to those in healthy-Tr recipients.

**Figure 2.**
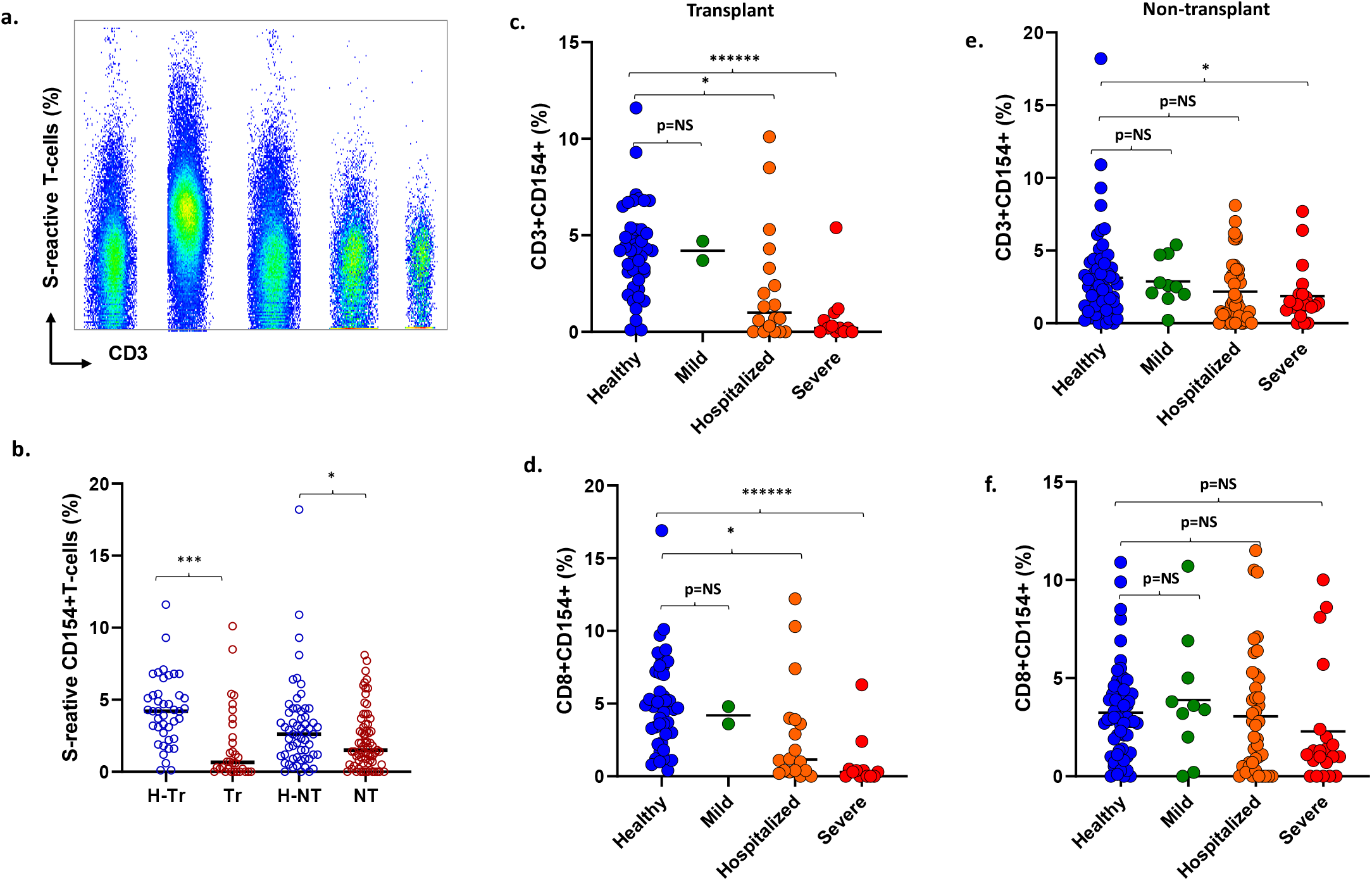
**a.** Flow cytometry scatterplots S-reactive CD3 cells in a representative healthy-transplant, healthy-non-transplant, Mild COVID-19, COVID-19 hospitalized, and COVID-19 subject intubated for mechanical ventilation. **b**. Dot plots show frequencies of S-reactive T-cells (CD3) in healthy-transplant (H-Tr), COVID-19-transplant (Tr), healthy-non-transplant (H-NT) and COVID-19 non-transplant (NT) subjects. **c**-**f**. Dot plots show frequencies of S-reactive CD3 cells (c, e) and CD8 cells (d, f) in transplant (c, d) and non-transplant patients (e, f) with COVID-19 who have mild infection treated as outpatient, or are hospitalized or have severe infection. Corresponding frequencies from healthy transplant and non-transplant subjects are shown in each dot plot (* represents p:value <0.05).

These differences were not seen in NT patients with COVID-19 compared with healthy-NT subjects. The sole exception consisted of lower S-reactive CD3 cells in NT patients with COVID-19, compared with healthy-NT subjects, p=0.045 (Figure 2e-f).

### Impaired antibody response to RBD in COVID-19 transplant patients

Of 74 COVID-19 patients with antibody measurements, 51 received convalescent plasma. IgG to spike antigen and RBD antigen were present in 49 of 51 (96%) and 47 of 51 (92%) patients, respectively. Among the remaining 23 patients who did not receive convalescent plasma, IgG to spike and RBD antigens were present in 21 (91%) and 16 (69.5%) patients, respectively. The incidence of anti-RBD IgG was significantly lower in transplant patients with COVID-19, 2 of 7 or 29%, compared with non-transplant patients, 14 of 16 or 88% (p=0.011) (Figure 3a). No differences were seen in the incidence of anti-spike IgG (5/7 or 71% vs 16/16 or 100%, p=NS) (Figure 3b). Subjects without and with anti-RBD antibody did not differ in timing of the sample from diagnosis (mean+/-SD 18+/-12.5 vs 12+/-12, p=0.258, NS, respectively), frequencies of S-reactive T-cells (mean 3.1+/-2.4% vs 1.8+/2%, p=0.225, NS, respectively), or proportions of patients requiring intubation (2/7 or 29% vs 4/16 or 25%, p=1.00, NS, respectively. S-reactive B-cell frequencies were also significantly lower in Tr and NT patients with COVID-19, compared with corresponding healthy subjects (Figure 3c).

**Figure 3.**
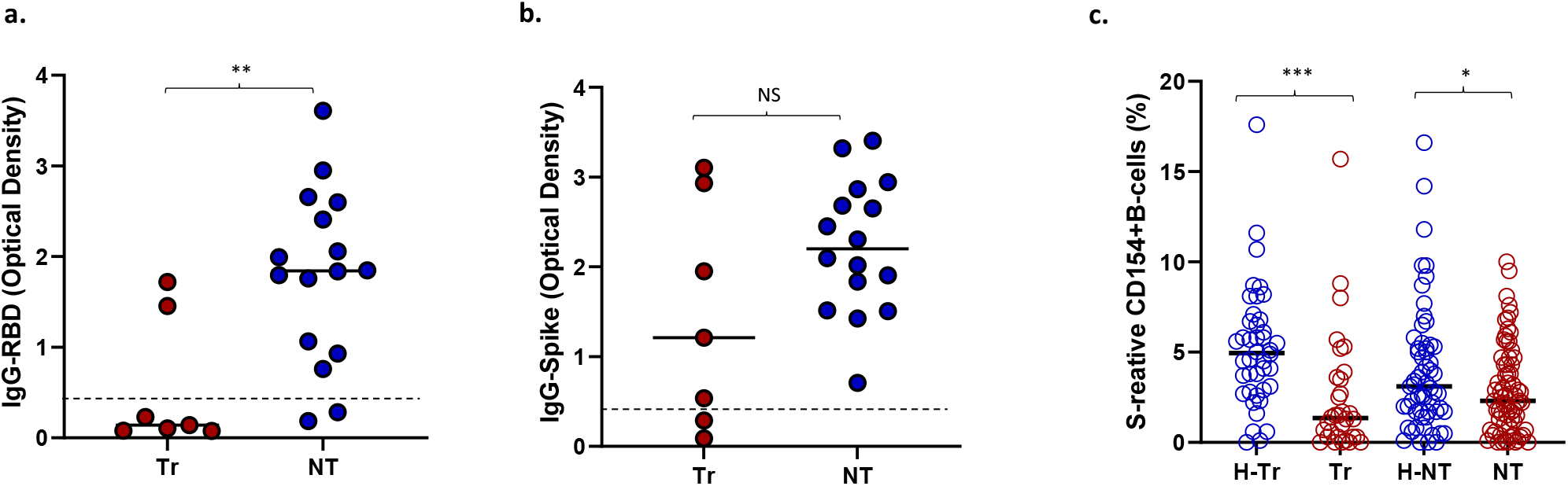
Optical density at 490 nm (OD_490_) for **(a)** Anti-RBD IgG and **(b)** Anti-spike IgG in transplant (Tr) and non-transplant (NT) patients with COVID-19 infection. Dotted lines show the OD_490_ cutoff of 0.45 above which the tests are deemed positive. **(c)** S-reactive B-cell frequencies in healthy-NT, healthy-T and T and NT patients with COVID-19 Infection. (* represents p:value <0.05).

### Increased risk of CMV co-infection in transplant recipients

Of 32 Tr recipients with COVID-19, 11 (34%) experienced CMV infection-10 had CMV viremia and one had CMV hepatitis. Consistent with this increased risk in Tr patients, CMV infection was associated with decreased T-cell immunity to this virus in Tr patients with COVID-19. Frequencies of CMV-specific T-cells which express CD154 after stimulation with the pp65 antigenic peptide mixture were measured as described previously, in 61 subjects^18^. CMV-specific T-cell frequencies were significantly lower in 16 Tr recipients with COVID-19 compared with 13 healthy Tr recipients (0.5+/-0.4% vs 1.5+/-0.5%, p=3E-05, Figure 4a). CMV-specific T-cell frequencies were not significantly different between 6 NT subjects without and 26 NT subjects with COVID-19 (p=0.21, NS, Figure 4b). CMV infection did not occur in NT patients with COVID-19.

**Figure 4.**
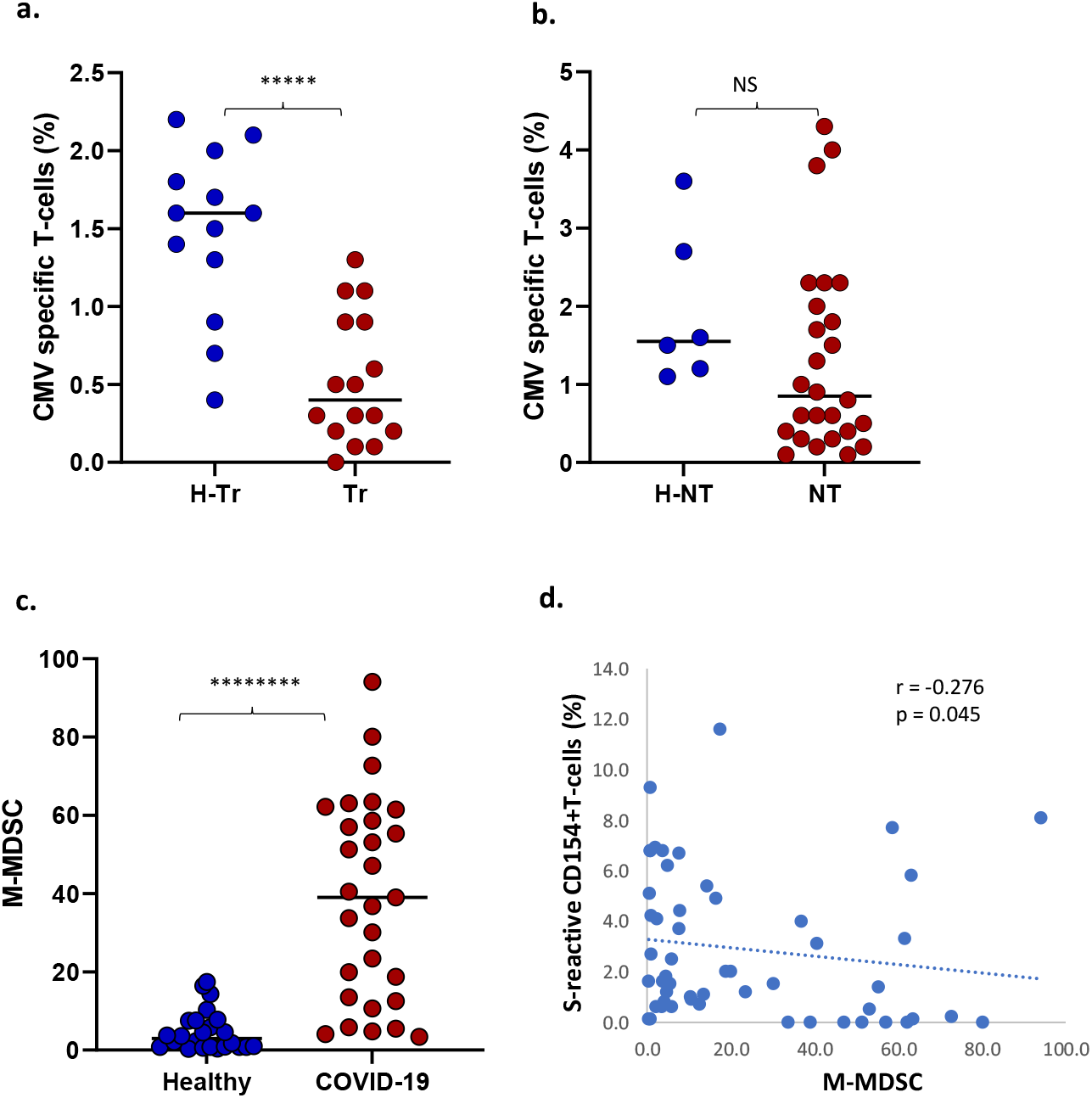
**a.** Dot plot shows CMV-specific T-cell frequencies among healthy transplant (H-Tr), healthy non-transplant (HNT), and COVID-19 patients with transplant (Tr) and without transplant (NT). **b**. frequencies of monocytic myeloid-derived suppressor cells (MDSC) in healthy unexposed subjects and COVID-19 patients. **c**. Correlation between frequencies of S-reactive CD154+T-cells and monocytic MDSC. (* represents p:value <0.05).

### Increased circulating myeloid-derived suppressor cells (MDSC) during COVID-19 infection

Twenty-four healthy and 29 COVID-19 patients were tested for circulating MDSCs. COVID-19 patients demonstrated higher frequencies of monocytic or M-MDSC (CD14^+^HLA-DR^-^) compared with healthy subjects (Median ±SEM, 39 ±7.8% vs 2.95 ±1.1%, p= 9.8 E-08) (Figure 4c). M-MDSC frequencies correlated negatively with S-reactive T-cell frequencies (Spearman’s r = −0.276, p= 0.045 Figure 4d). Polymorphonuclear or P-MDSC (CD15+CD14-CD11b+) frequencies were also higher in four COVID-19 subjects compared with 22 healthy subjects (median ±SEM, 64.2 ± 19.4% vs 1.25 ±1.4%, p= 0.059, NS), and were negatively correlated with S-reactive T-cell frequencies (Spearman’s r = −0.518, p= 0.007).

## Discussion

Our study found that S-reactive T-cells are present in pre-pandemic PBL samples from chronically immunosuppressed transplanted (Tr) recipients. This type of pre-existing T-cell immunity has been reported previously in the general population and is also seen in our study population of healthy NT subjects^12-15^. Experimental evidence from previous studies implicates prior exposure to structurally similar human coronaviruses, which cause seasonal flu^16-17^. We speculate that this explanation also applies to our Tr recipient cohort. Unlike some previous studies, however, we observed lower S-reactive T-cell frequencies in COVID-19 patients compared with healthy unexposed individuals. This decrease was significant and most pronounced for Tr patients with COVID-19 compared with controls (Figure 2b-d). Further, compared with healthy-Tr subjects, Tr patients with COVID-19 infection also demonstrated a progressive decline in S-reactive T-cell frequencies with increasing disease severity, from hospitalization (p<0.05) to severe disease requiring intubation (p<0.0001). S-reactive T-cell frequencies in Tr patients with mild symptoms were in the same range as healthy unexposed Tr subjects (p=NS). These differences were also observed for S-reactive CD8 cells among Tr patients with COVID-19 (Figure 2d). Among NT patients with COVID-19, the decrease in S-reactive CD3 cell frequencies arose from those with severe COVID-19 infection (Figure 2e). No other between-group differences were identified for NT subjects.

Unlike previous studies, the majority of our COVID-19 patients, 91 of 103, were hospitalized, 58 without and 33 with severe disease requiring intubation for respiratory failure. This distribution represents a more severely affected infection cohort and may explain lower mean T-cell frequencies in infected patients compared with those who were healthy. Loss of T-cell immunity to the virus has been observed in critically ill patients in some previous studies^12^. Previous reports have also shown higher S-reactive T-cell frequencies in convalescent patients compared with unexposed subjects^12-15^. These higher responses may be unique to the convalescent phase. Another reason for the higher T-cell responses in COVID-19 infection in some previous studies may be the use of peptides with high affinity for selected HLA specificities, with or without adjunctive co-stimulators. This approach may have elicited larger T-cell responses from memory subsets. Our study patients were sampled at an average interval of 12 days after diagnosis of COVID-19 and assayed using unselected peptide stimulators, without adjunctive co-stimulators.

In previous studies, S-reactive T-cell frequencies averaging <1% have been observed in healthy unexposed subjects, compared with roughly 3% in our studies^12-15^. Some of these studies counted S-reactive T-cells as those that co-expressed marker combinations like CD137 and CD69, but excluded S-reactive T-cells that expressed either marker alone^12,13^. We have modeled our assay on clinical assays which measure antiviral T-cell immunity by employing a single marker. These assays use either IFNγ or CD154 as a marker of antigen-specific T-cells ^18-20,22,23^. CD154 can substitute for IFNγ because it is co-expressed in viral antigen-specific T-cells^18,24^. As a single marker of antigen-specific T-cells, CD154 can also be detected with non-permeabilizing methods. Alloantigen-specific T-cytotoxic memory cells that express CD154 have met criteria for regulatory approval to predict transplant rejection^25^. S-reactive T-cell frequencies averaging 3% in our healthy unexposed subjects have also been observed among proliferating S-reactive T-cells in a previous study^14^. We cannot fully explain higher average frequencies in healthy unexposed Tr compared with NT subjects (mean 3.1 vs 4.2 %, p=0.042, Table S6). However, extended ex vivo exposure of normal human PBL to pro-apoptotic anti-lymphocyte antibodies enriches apoptosis-resistant alloantigen-reactive CD154+T-cells among surviving PBL^26^. Thus, it is possible that exposure of T-cells to chronic immunosuppression may have contributed to an enrichment of S-reactive T-cells in PBL from Tr recipients.

The Tr recipient cohort with COVID-19 was noteworthy for CMV co-infection presenting as viremia in 10, and CMV hepatitis in one recipient for an incidence of 11 of 32 or 34%. CMV infection occurred at a median of 22 days (range 1-104 days) after diagnosis of COVID-19 infection. Transplant recipients with COVID-19 also demonstrated lower frequencies of CMV-specific T-cells compared with NT COVID-19 patients, 0.4 ±0.1 vs 0.85 ±0.24, p=0.0048. Consistent with a lack of such differences in NT subjects, no CMV co-infections were reported in NT patients with COVID-19.

Of great interest is the observation that Tr recipients also demonstrated a lower incidence of IgG antibodies to the RBD component of the S protein after COVID-19 infection compared with NT recipients, 2 of 7 vs 3 of 16, p=0.011. The incidence of anti-S IgG antibodies was similar between the T and NT groups. The RBD sequence is a component of the less conserved N-terminal S1 sequence of the SARS-CoV-2 spike protein. The S1 protein has 60% sequence similarity to hCoV. As such, the RBD sequence may be less immunogenic when presented to the host immune system for the first time, compared with the more conserved C-terminal S2 sequence, which has 80% homology with hCoV. Test positivity was based on an OD_490_ of 0.45 or greater in the ELISA antibody binding assay. The amount of IgG antibody reflected by OD_490_ readings was also lower in Tr compared with NT patients for anti-spike IgG (p=0.16, NS) achieving significance for anti-RBD IgG (p<0.001) (Figure 3). Impaired antibody responses to natural COVID-19 infection in Tr recipients may augur impaired antibody responses to COVID-19 vaccines in this population.

Suppressed cellular and antibody responses in Tr recipients may have other reasons. Recent studies have revealed increased circulating myeloid derived suppressor cells (MDSC), pyroptotic cell death and lymphopenia in COVID-19 patients^27-31^. MDSC are myeloid progenitors that expand in peripheral blood in response to lymphopenia and are known to suppress T-cells. Frequencies of monocytic and polymorphonuclear MDSC were higher in COVID-19 patients compared with healthy unexposed subjects. The corresponding decrease in S-reactive T-cells is reflected in significant negative correlations between S-reactive T-cells and MDSC.

In conclusion, transplant recipients demonstrate pre-existing T-cell immunity to SARS-CoV-2 in a manner similar to the general population. Unique attributes of COVID-19 infection in transplant recipients include a) impaired T-cell immunity to SARS-CoV-2, to the greatest degree in those with increasing disease-severity, b) increased risk for CMV co-infection, and c) impaired antibody responses. Surveillance of CMV viral loads during COVID-19 infection, and post-vaccination surveillance of antibody responses to confirm vaccine efficacy may be necessary in transplant recipients.

## Methods

Human Subjects: COVID-19 patients were enrolled under IRB-approved protocols 2017-0365, Pro00101915, and 1551551 respectively, at three centers in Washington, DC, Charleston, SC, and Edinburg, TX, respectively. De-identified residual cryopreserved PBL samples were tested under IRB-exempt protocol, and samples from healthy-NT subjects were tested under IRB approved protocol 6774 in the reference laboratory (Plexision, Pittsburgh, PA). Healthy unexposed subjects, H-NT and H-Tr were tested using samples that were either obtained pre-pandemic, in 2019 or earlier, or were tested after confirming absence of symptoms suggestive of flu-like symptoms in the 6-month period prior to testing and a negative test for IgG to S and RBD antigens. COVID-19 patients, Tr or NT, were tested with samples obtained after confirmation of diagnosis with PCR.

### Measuring SARS-CoV-2-reactive T-cell and B-cell subsets

All PBL samples were cultured alone (background), with 315 15-mer overlapping peptides with 11-mer overlap representing the 1273 amino acid spike antigen (test reaction), and with phorbol-myristic acid-Calcium ionophor (PMA, positive control) for 16 hours at 37°C in 5% CO2 incubator. The peptide mixture consisted of two components mixed in equal parts-158 peptides representing the less conserved N-terminal sequence, S1, and 157 peptides representing the more conserved C-terminal sequence, S2, of the spike protein (JPT Peptides, Berlin, Germany). The S1 and S2 sequences respectively have 64% and 90% sequence homology with the SARS virus^32^. The culture medium contained fluorochrome-labeled antibody to CD154 (catalog #563886, BD Biosciences, San Jose, CA). Cells were acquired on the FACS-Canto II flow cytometer with blue, red and violet lasers after addition of fluorochrome labeled antibodies to CD3, CD4, CD8, and CD19 and the viability dye 7-aminoactinomycin-D (catalog #s 340662, 641407, 340692, 341103, 559925, respectively, BD Biosciences, San Jose, CA). The gating strategy is shown in Figure S1. Scatterplots acquired from assay reaction conditions for CD3, CD4, CD8 and CD19 cells are shown in Figure S2. Frequencies for each subset which were reactive to the S peptide mixture were analyzed further after subtracting corresponding background frequencies.

### CMV- and mitogen-reactive T-cells

Previously described methods were used to measure frequencies of CMV-specific T-cells and mitogen-reactive T-cells that expressed CD154 in response to stimulation with the pp65-CMV antigen and PMA, respectively^18^.

### Serological assay to detect SARS-CoV-2 antibody

96-well microtiter plates were coated overnight at 4°C with commercially available S-protein (Cat # 46328, LakePharma, San Carlos, CA,) at 2 ug/ml, and blocked for 1hr with PBS-Tween + 3% milk powder (weight/volume). Precoated wells were incubated with diluted samples for 2 hours, followed by anti-human IgG (Fab specific) HRP labeled secondary antibody 1:3000 in PBS-T containing 1% milk for 1 hour. After adding substrate (OPD solution), followed by 50 μl of 3M hydrochloric acid to stop the reaction, plates were read at 490 nm on a spectrophotometer. With all samples, inactivated human AB serum was used as a negative control, while monoclonal antibody CR3022 was used as a positive control. Results were read on a plate reader as optical density at 490 nm. An optical density of 0.45 or greater was considered a positive test as reported earlier^33^.

### Myeloid-derived suppressor cells (MDSC)

MDSC represent early lineage cells that cause T-cell suppression and develop in response to lymphopenia and the inflammatory response to the viral infection^34-36^. Fluorochrome-labeled antibodies to the respective markers for each cell were used to characterize monocytic- and polymorphonuclear-MDSC (M-MDSC and P-MDSC). The respective phenotypes were CD14+HLADR- and CD15+CD14-CD11b+^36^. Antibodies used were from Biolegend (Cat # 307618,301906,301306, San Diego, CA) or BD Biosciences (Cat # 563743, San Jose, CA).

### Statistical methods

Descriptive statistics were used to summarize group features. Between group comparisons were performed with t-tests for unadjusted data and linear models to adjust for demographic variables.

## Acknowledgements

NSF#2033307 (CA, Plexision), NIH Grant number UL1 TR001450 (MUSC), intramural support from all participating institutions and Plexision.

## Data Availability Statement

All data that underlie the results reported in this Article (including study protocol) on individual participants will be made available to researchers who provide a methodologically sound proposal to the corresponding author.

## Contributors and Data Management

RV, AHK, JA, GM, SN, SR, HD, KK, MMBC, KS, GB, AK, MN, PB, TF HD, KK recruited subjects, interpreted results, wrote and edited manuscript. SN and SM performed, interpreted and described antibody testing. CA performed and described CMI assays for SARS2 antigens on de-identified samples. BS, MS, NA and ES compiled, cross-checked, tabulated, summarized and described cell counts and frequency data. Demographics of de-identified subjects were summarized by BS who also performed and described CMI assays for CMV. Cytometry results and demographics were verified by PS, and transmitted to statistician BWH who merged the two datasets, performed all analyses and returned results and descriptions of analyses to PS and wrote and edited manuscript. KM and SS confirmed results of logistic regression with alternative linear models, edited and wrote manuscript. PS interpreted study results and relayed interpretations to RS for communication to all investigators. RS conceived the study, coordinated with centers and investigators, incorporated descriptions from other authors, wrote and edited manuscript with all authors.

## Abbreviations

(CMI): cell-mediated immunity
(SARS2): SARS-CoV-2
(H-NT): healthy-non transplant
(H-Tr): healthy transplant
(Tr): COVID-19 transplant
(NT): COVID-19 non-transplant
(TP): true positive
(FN): false negative
(PBL): peripheral blood leukocytes
(M-MDSC and P-MDSC): monocytic- and polymorphonuclear-MDSC

## Supplementary Figure Legends

**Figure S1.**
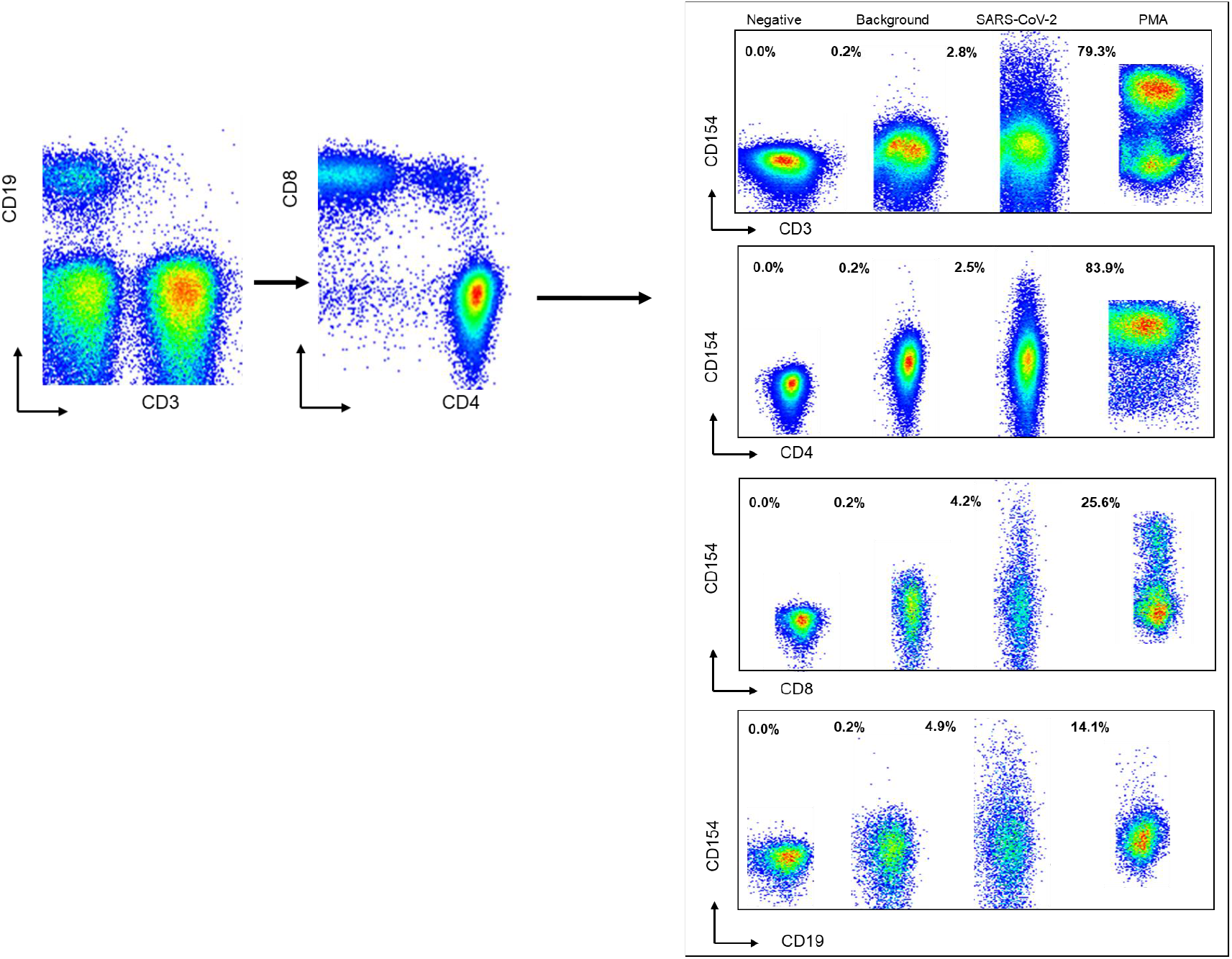
Flow cytometric gating strategy shows derivation of CD3^+^ T-cells and CD19^+^ B-cells from the lymphocyte population. CD4 and CD8 T-cells were than gated from CD3^+^ T-cells. Scatterplots show CD3, CD4, CD8 and CD19 cells that express CD154 when incubated alone (background), with spike antigen (test reaction) and PMA-Calcium ionophore (positive control). The negative control reaction shows autofluorescence in the absence of fluorochrome-labeled CD154 antibody.

**Table S1:** Demographics, treatment and outcomes of 66 patients with COVID-19 infection. Type (Tr= Transplant, NT= No-Transplant), Race (C= Caucasian, AA=African American, H=Hispanic, A=Asian), Status (A= Alive, D=Dead), Plasma treatment (N=No, Y=Yes), Dexamethasone / Prednisone. Dexamethasone is given as part of the COVID-19 treatment regime, Prednisone is given to transplant patients as part of maintenance immunosuppression.

| Type | Race | Age | Gender | Days from diagnosis | Disease Severity | Status | Plasma treatment | Dexamethasone / Prednisone |
| --- | --- | --- | --- | --- | --- | --- | --- | --- |
| NT | C | 47 | F | 18 | Intubated | A | Y | Dexamethasone |
| NT | H | 61 | M | 14 | Intubated | A | Y | Dexamethasone |
| NT | H | 63 | F | 26 | Intubated | A | Y | Dexamethasone |
| NT | H | 44 | F | 21 | Intubated | A | Y | Dexamethasone |
| NT | H | 59 | M | 0 | Intubated | A | Y | Dexamethasone |
| NT | H | 48 | F | 22 | Intubated | D | Y | Dexamethasone |
| NT | H | 69 | M | 6 | Intubated | D | Y | Dexamethasone |
| NT | H | 74 | M | 88 | Intubated | A | Y | Dexamethasone |
| NT | H | 71 | M | 4 | Intubated | A | Y | Dexamethasone |
| NT | H | 61 | M | 18 | Intubated | A | Y | Dexamethasone |
| NT | H | 67 | M | 17 | Intubated | A | Y | None |
| NT | H | 57 | M | 14 | Intubated | A | Y | N |
| NT | H | 45 | M | 15 | Intubated | A | Y | N |
| NT | H | 53 | M | 29 | Intubated | A | Y | N |
| NT | H | 58 | F | 26 | Intubated | A | Y | N |
| NT | H | 70 | F | 16 | Intubated | A | Y | N |
| NT | H | 70 | M | 19 | Intubated | D | Y | N |
| NT | AA | 24 | F | 2 | Intubated | D | N | N |
| NT | H | 57 | F | 14 | Intubated | A | Y | N |
| NT | H | 77 | F | 4 | Intubated | A | Y | N |
| NT | H | 54 | F | 5 | Intubated | A | Y | N |
| NT | H | 30 | M | 1 | Hospitalized | A | N | **Dexamethasone** |
| NT | H | 47 | M | 30 | Hospitalized | A | N | **Dexamethasone** |
| NT | H | 32 | M | 3 | Hospitalized | A | Y | **Dexamethasone** |
| NT | H | 34 | F | 11 | Hospitalized | A | Y | **Dexamethasone** |
| NT | H | 44 | F | 2 | Hospitalized | A | Y | **Dexamethasone** |
| NT | H | 56 | M | 5 | Hospitalized | A | Y | **Dexamethasone** |
| NT | H | 24 | M | 7 | Hospitalized | A | Y | **Dexamethasone** |
| NT | H | 55 | M | 4 | Hospitalized | A | Y | **Dexamethasone** |
| NT | H | 35 | F | 11 | Hospitalized | A | Y | **Dexamethasone** |
| NT | H | 51 | M | 8 | Hospitalized | A | Y | **Dexamethasone** |
| NT | H | 68 | M | 3 | Hospitalized | A | Y | **Dexamethasone** |
| NT | H | 65 | F | 5 | Hospitalized | A | Y | **Dexamethasone** |
| NT | H | 87 | F | 6 | Hospitalized | A | Y | **Dexamethasone** |
| NT | H | 35 | F | 0 | Hospitalized | A | Y | **Dexamethasone** |
| NT | H | 70 | M | 1 | Hospitalized | A | Y | **Dexamethasone** |
| NT | H | 79 | M | 0 | Hospitalized | A | Y | **Dexamethasone** |
| NT | H | 83 | M | 0 | Hospitalized | A | Y | **Dexamethasone** |
| NT | H | 72 | M | 0 | Hospitalized | A | Y | **Dexamethasone** |
| NT | H | 75 | F | 0 | Hospitalized | A | Y | **Dexamethasone** |
| NT | H | 69 | M | 0 | Hospitalized | A | Y | **Dexamethasone** |
| NT | H | 50 | M | 0 | Hospitalized | A | Y | **Dexamethasone** |
| NT | C | 51 | F | 94 | Hospitalized | A | Y | **Dexamethasone** |
| NT | H | 74 | M | 3 | Hospitalized | A | Y | **Dexamethasone** |
| NT | H | 65 | M | 27 | Hospitalized | A | Y | **Dexamethasone** |
| NT | H | 87 | F | 8 | Hospitalized | A | Y | **Dexamethasone** |
| NT | C | 72 | F | 4 | Hospitalized | A | Y | **Dexamethasone** |
| NT | H | 65 | F | 1 | Hospitalized | A | Y | **Dexamethasone** |
| NT | H | 73 | M | 2 | Hospitalized | A | Y | **Dexamethasone** |
| NT | H | 49 | F | 2 | Hospitalized | A | N | None |
| NT | H | 78 | M | 8 | Hospitalized | A | N | None |
| NT | H | 65 | M | 1 | Hospitalized | A | Y | None |
| NT | H | 33 | M | 28 | Hospitalized | A | Y | None |
| NT | H | 52 | M | 27 | Hospitalized | A | Y | None |
| NT | AA | 63 | F | 2 | Hospitalized | A | N | None |
| NT | AA | 69 | M | 2 | Hospitalized | A | N | None |
| NT | H | 78 | F | 3 | Hospitalized | A | N | None |
| NT | H | 66 | F | 1 | Hospitalized | A | N | None |
| NT | H | 69 | M | 0 | Hospitalized | A | N | None |
| NT | H | 53 | F | 9 | Hospitalized | A | Y | None |
| NT | AA | 65 | M | 2 | Hospitalized | A | N | **Dexamethasone** |
| NT | C | 46 | M | 18 | Mild/Asymp | A | N | None |
| NT | A | 32 | M | 16 | Mild/Asymp | A | N | None |
| NT | C | 36 | M | 14 | Mild/Asymp | A | N | None |
| NT | C | 26 | F | 14 | Mild/Asymp | A | N | None |
| NT | C | 39 | M | 16 | Mild/Asymp | A | N | None |
| NT | C | 25 | F | 18 | Mild/Asymp | A | N | None |
| NT | C | 28 | F | 16 | Mild/Asymp | A | N | None |
| NT | C | 25 | F | 24 | Mild/Asymp | A | N | None |
| NT | C | 51 | F | 26 | Mild/Asymp | A | N | None |
| NT | C | 38 | F | 21 | Mild/Asymp | A | N | None |
| Tr | AA | 53 | M | 2 | Intubated | A | N | Prednisone |
| Tr | H | 76 | M | 14 | Intubated | A | y | Prednisone |
| Tr | H | 58 | M | 29 | Intubated | A | N | Prednisone |
| Tr | H | 45 | M | 4 | Intubated | A | N | Prednisone |
| Tr | AA | 43 | F | 4 | Intubated | A | N | Prednisone |
| Tr | C | 75 | F | 4 | Intubated | A | N | Prednisone |
| Tr | C | 63 | M | 26 | Intubated | A | N | Prednisone |
| Tr | C | 68 | M | 7 | Intubated | D | N | Prednisone |
| Tr | AA | 44 | F | 19 | Intubated | D | N | Prednisone |
| Tr | C | 73 | M | 23 | Intubated | D | N | Prednisone |
| Tr | C | 46 | M | 43 | Intubated | D | N | Prednisone |
| Tr | H | 62 | M | 17 | Intubated | D | Y | Prednisone |
| Tr | AA | 72 | M | 26 | Hospitalized | A | N | Prednisone |
| Tr | H | 75 | M | 5 | Hospitalized | A | N | Prednisone |
| Tr | AA | 77 | F | 11 | Hospitalized | A | N | Prednisone |
| Tr | C | 41 | M | 2 | Hospitalized | A | N | Prednisone |
| Tr | H | 15 | M | 3 | Hospitalized | A | N | Prednisone |
| Tr | H | 67 | F | 14 | Hospitalized | A | N | Prednisone |
| Tr | C | 75 | F | 9 | Hospitalized | A | N | Prednisone |
| Tr | AA | 33 | F | 5 | Hospitalized | A | N | Prednisone |
| Tr | AA | 44 | M | 5 | Hospitalized | A | N | Prednisone |
| Tr | AA | 64 | F | 2 | Hospitalized | A | N | Prednisone |
| Tr | C | 51 | F | 2 | Hospitalized | A | N | Prednisone |
| Tr | AA | 43 | M | 2 | Hospitalized | A | N | Prednisone |
| Tr | H | 39 | M | 11 | Hospitalized | A | N | Prednisone |
| Tr | AA | 68 | M | 32 | Hospitalized | A | N | Prednisone |
| Tr | AA | 31 | M | 6 | Hospitalized | A | N | Prednisone |
| Tr | **A** | 1 | M | 3 | Hospitalized | A | N | None |
| Tr | H | 51 | F | 5 | Hospitalized | A | Y | Prednisone |
| Tr | H | 7 | F | 3 | Hospitalized | A | Y | None |
| Tr | H | 1 | M | 50 | Mild/Asymp | A | N | Prednisone |
| Tr | A | 17 | M | 30 | Mild/Asymp | A | N | Prednisone |

## SUPPLEMENTARY TABLES

**Table S2:** Summary of variation in CD154+PBL subsets (CD3, CD4, CD8, CD19) measured in same-day duplicate testing of five PBL samples in response to spike protein (upper half of table) and PMA stimulation (lower half of table). Variation is measured as the coefficient of variation (CV %).

| Spike | N | Mean CV | CI low | CI-up | SD | Median | Min | Max |
| --- | --- | --- | --- | --- | --- | --- | --- | --- |
| CD3 | 5.0 | 3.0 | -0.8 | 6.8 | 4.3 | 1.9 | 0.0 | 10.4 |
| CD4 | 5.0 | 5.0 | -0.5 | 10.5 | 6.3 | 2.2 | 0.0 | 14.6 |
| CD8 | 5.0 | 3.3 | -0.2 | 6.8 | 3.9 | 1.8 | 0.0 | 10.0 |
| CD19 | 5.0 | 2.0 | 0.9 | 3.2 | 1.3 | 2.0 | 0.0 | 3.6 |
| PMA | N | Mean CV | CI low | CI-up | SD | Median | Min | Max |
| CD3 | 5.0 | 3.6 | 2.1 | 5.1 | 1.7 | 4.4 | 1.1 | 5.0 |
| CD4 | 5.0 | 2.6 | -0.5 | 5.8 | 3.6 | 2.0 | 0.1 | 8.8 |
| CD8 | 5.0 | 3.5 | 1.9 | 5.1 | 1.8 | 3.7 | 1.3 | 6.2 |
| CD19 | 5.0 | 3.0 | 0.8 | 5.2 | 2.5 | 2.0 | 0.8 | 6.7 |

**Table S3:**
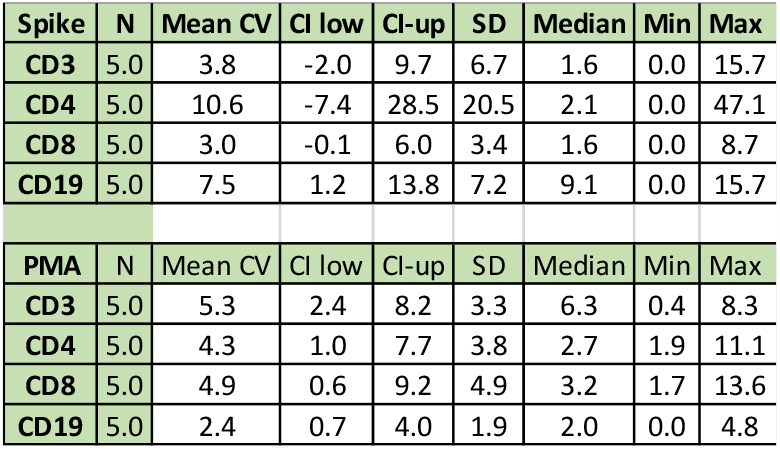
Summary of variation in CD154+PBL subsets (CD3, CD4, CD8, CD19) measured in five PBL samples tested on the day of phlebotomy and after cryopreservation for 7 days. All samples were stimulated with spike protein (upper half of table) and PMA (lower half of table). Variation is measured as the coefficient of variation (CV %).

| Spike | N | Mean CV | CI low | CI-up | SD | Median | Min | Max |
| --- | --- | --- | --- | --- | --- | --- | --- | --- |
| CD3 | 5.0 | 3.8 | -2.0 | 9.7 | 6.7 | 1.6 | 0.0 | 15.7 |
| CD4 | 5.0 | 10.6 | -7.4 | 28.5 | 20.5 | 2.1 | 0.0 | 47.1 |
| CD8 | 5.0 | 3.0 | -0.1 | 6.0 | 3.4 | 1.6 | 0.0 | 8.7 |
| CD19 | 5.0 | 7.5 | 1.2 | 13.8 | 7.2 | 9.1 | 0.0 | 15.7 |
| PMA | N | Mean CV | CI low | CI-up | SD | Median | Min | Max |
| CD3 | 5.0 | 5.3 | 2.4 | 8.2 | 3.3 | 6.3 | 0.4 | 8.3 |
| CD4 | 5.0 | 4.3 | 1.0 | 7.7 | 3.8 | 2.7 | 1.9 | 11.1 |
| CD8 | 5.0 | 4.9 | 0.6 | 9.2 | 4.9 | 3.2 | 1.7 | 13.6 |
| CD19 | 5.0 | 2.4 | 0.7 | 4.0 | 1.9 | 2.0 | 0.0 | 4.8 |

**Table S4.** Summary of variation in CD154+PBL subsets (CD3, CD4, CD8, CD19) measured in five PBL samples tested on the day of phlebotomy and after overnight storage at room temperature. All samples were stimulated with spike protein (upper half of table) and PMA (lower half of table). Variation is measured as the coefficient of variation (CV %).

| Spike | N | Mean CV | CI low | CI-up | SD | Median | Min | Max |
| --- | --- | --- | --- | --- | --- | --- | --- | --- |
| CD3 | 5.0 | 6.3 | 0.1 | 12.6 | 7.1 | 4.0 | 0.0 | 15.7 |
| CD4 | 5.0 | 7.8 | 1.3 | 14.3 | 7.4 | 2.8 | 2.1 | 16.1 |
| CD8 | 5.0 | 6.6 | 0.6 | 12.5 | 6.8 | 1.9 | 1.2 | 15.1 |
| CD19 | 5.0 | 6.5 | 2.8 | 10.2 | 4.2 | 4.7 | 2.8 | 13.1 |
| PMA | N | Mean CV | CI low | CI-up | SD | Median | Min | Max |
| CD3 | 5.0 | 3.9 | 0.4 | 7.4 | 4.0 | 2.5 | 0.4 | 10.7 |
| CD4 | 5.0 | 3.4 | 0.9 | 5.9 | 2.9 | 3.4 | 0.7 | 7.8 |
| CD8 | 5.0 | 6.2 | 1.4 | 11.1 | 5.5 | 5.6 | 1.1 | 15.5 |
| CD19 | 5.0 | 5.4 | 3.2 | 7.6 | 2.5 | 6.0 | 2.0 | 7.8 |

**Table S5.** Summary of variation in CD154+PBL subsets (CD3, CD4, CD8, CD19) measured in five PBL samples tested on the day of phlebotomy and after overnight shipment at ambient temperature. All samples were stimulated with spike protein (upper half of table) and PMA (lower half of table). Variation is measured as the coefficient of variation (CV %).

| Spike | N | Mean CV | CI low | CI-up | SD | Median | Min | Max |
| --- | --- | --- | --- | --- | --- | --- | --- | --- |
| CD3 | 5.0 | 7.2 | 1.9 | 12.5 | 6.0 | 6.0 | 0.0 | 15.7 |
| CD4 | 5.0 | 9.3 | 2.2 | 16.4 | 8.1 | 4.4 | 2.1 | 20.2 |
| CD8 | 5.0 | 4.0 | 0.2 | 7.8 | 4.3 | 3.3 | 0.0 | 11.3 |
| CD19 | 5.0 | 4.6 | 1.3 | 8.0 | 3.8 | 2.8 | 1.0 | 9.1 |
| PMA | N | Mean CV | CI low | CI-up | SD | Median | Min | Max |
| CD3 | 5.0 | 4.1 | 0.8 | 7.5 | 3.8 | 1.9 | 1.4 | 10.4 |
| CD4 | 5.0 | 3.0 | 1.4 | 4.6 | 1.8 | 2.8 | 1.1 | 5.2 |
| CD8 | 5.0 | 5.7 | 3.9 | 7.5 | 2.0 | 5.3 | 3.2 | 8.8 |
| CD19 | 5.0 | 3.2 | 0.1 | 6.3 | 3.5 | 1.5 | 0.0 | 7.2 |

**Table S6.** Summary data for mean and median frequencies of S-reactive CD154+PBL subsets.

|  |  | **CD3** | **CD4** | **CD8** | **CD19** |
| --- | --- | --- | --- | --- | --- |
| **Healthy-NT**<br>**(N=59)** | Mean | 3.1 | 2.8 | 3.2 | 4.0 |
|  | Median | 2.6 | 2.3 | 3.0 | 3.1 |
|  | SD | 3.0 | 2.5 | 2.3 | 3.5 |
| **Healthy-Tr**<br>**(N=42)** | Mean | 4.2 | 4.0 | 5.0 | 5.2 |
|  | Median | 4.2 | 3.8 | 4.8 | 5.0 |
|  | SD | 2.4 | 2.2 | 3.2 | 3.3 |
| **COVID-19-NT**<br>**(N=71)** | Mean | 2.2 | 1.9 | 2.9 | 2.9 |
|  | Median | 1.5 | 1.3 | 1.9 | 2.3 |
|  | SD | 2.1 | 1.8 | 3.1 | 2.4 |
| **COVID-19-Tr**<br>**(N=32)** | Mean | 1.8 | 1.8 | 2.2 | 2.5 |
|  | Median | 0.7 | 0.6 | 0.5 | 1.4 |
|  | SD | 2.6 | 2.5 | 3.1 | 3.4 |
| **p-value** | H-NT vs H-Tr | 0.0422 | 0.0147 | 0.0043 | 0.0755 |
|  | H-NT vs NT | 0.0437 | 0.0222 | 0.5310 | 0.0497 |
|  | H-Tr vs Tr | 0.0001 | 0.0002 | 0.0004 | 0.0010 |

